# Delivering genes across the blood-brain barrier: LY6A, a novel cellular receptor for AAV-PHP.B capsids

**DOI:** 10.1101/538421

**Authors:** Qin Huang, Ken Y. Chan, Isabelle G. Tobey, Yujia Alina Chan, Tim Poterba, Christine L. Boutros, Alejandro B. Balazs, Richard Daneman, Jonathan M. Bloom, Cotton Seed, Benjamin E. Deverman

## Abstract

The engineered AAV-PHP.B family of adeno-associated virus efficiently delivers genes throughout the mouse central nervous system. To guide their application across disease models, and to inspire the development of translational gene therapy vectors useful for targeting neurological diseases in humans, we sought to elucidate the host factors responsible for the CNS tropism of AAV-PHP.B vectors. Leveraging CNS tropism differences across mouse strains, we conducted a genome-wide association study, and rapidly identified and verified LY6A as an essential receptor for the AAV-PHP.B vectors in brain endothelial cells. Importantly, this newly discovered mode of AAV binding and transduction is independent of other known AAV receptors and can be imported into different cell types to confer enhanced transduction by the AAV-PHP.B vectors.

## Introduction

With the boom of gene replacement, knockdown, and editing technologies, the number of diseases that are potentially treatable by gene therapy is rapidly expanding. AAV vectors are proving to be safe, versatile vehicles for *in vivo* gene therapy applications^1–4^. However, delivery challenges impede the application of gene therapy, particularly in the context of the brain, which is protected by the blood-brain barrier (BBB). To improve gene delivery across the central nervous system (CNS), our group and others have engineered AAV capsids using *in vivo* selection and directed evolution^5–8^. Our group has focused on engineering AAV9 variants, such as AAV-PHP.B^6^ and its further evolved, more efficient variant, AAV-PHP.eB^5^, to cross the adult BBB and enable efficient gene transfer to the CNS. Since their invention, AAV-PHP.B and AAV-PHP.eB have been applied across a wide range of neuroscience experiments in mice^5,9,10^, including genetic deficit correction^11,12^ and neurological disease modeling^13^.

A critical question has been how the AAV-PHP.B vectors cross the BBB and whether this mechanism can be translated to other species and, ultimately, humans. The enhanced CNS tropism of AAV-PHP.B and AAV-PHP.eB appears to extend to rats^14,15^, whereas studies testing AAV-PHP.B or related capsids in nonhuman primates (NHPs) have yielded differing outcomes^16–18^. Surprisingly, the enhanced CNS tropism of AAV-PHP.B^6,9,12–15,19,20^ is starkly absent in BALB/cJ mice^16^. These findings indicate that the ability of AAV-PHP.B to cross the BBB is affected by genetic factors that vary by species and mouse strain. In this work, we leverage this strain-dependence to identify LY6A as the cellular receptor responsible for the enhanced CNS tropism exhibited by the AAV-PHP.B capsid family. We demonstrate that LY6A-mediated transduction is independent of known AAV9 receptors and is a unique means by which AAV-PHP.B capsids cross the mouse BBB. This has widespread implications for guiding the use of AAV-PHP.B capsids in disease models, as well as ongoing efforts to engineer next-generation AAVs that cross the BBB in other species.

## Results

### *Ly6* genetic variants associate with the CNS tropism of AAV-PHP.eB

The dramatic difference in the CNS tropism of AAV-PHP.B in C57BL/6J versus BALB/cJ mice^16^ extended to AAV-PHP.eB (Fig. 1A) and two other AAV-PHP.B capsids (AAV-PHP.B2 and AAV-PHP.B3; Supplementary Fig. 1). Consistent with the reduced transduction phenotype in BALB/cJ mice, we observed a loss of the CNS-specific enhanced accumulation of AAV-PHP.eB, relative to AAV9, along the vasculature and within the brains of BALB/cJ mice (Fig. 1B, C). These findings suggest that a cellular factor present on the brain endothelium is responsible for the efficient CNS-wide transduction mediated by the AAV-PHP.B capsids, and that this factor is absent or nonfunctional in BALB/cJ mice.

**Figure 1.**
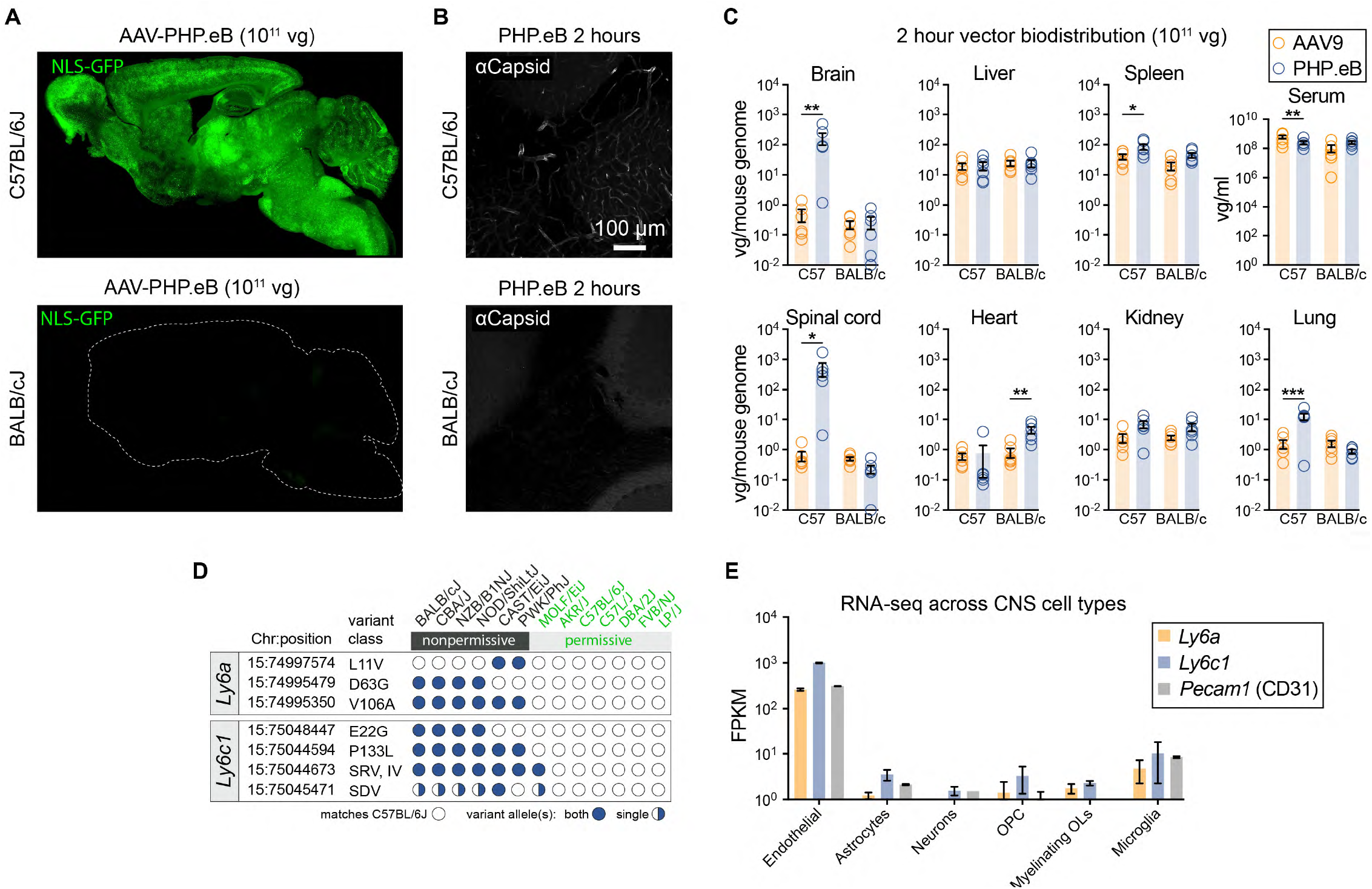
The nonpermissive AAV-PHP.eB CNS tropism phenotype associates with *Ly6* SNPs. (**A**) GFP within sagittal brain sections from C57BL/6J or BALB/cJ mice two weeks after intravenous administration of AAV-PHP.eB:CAG-NLS-GFP. (**B**) AAV capsid IHC within the cerebellum one hour after intravenous injection of AAV-PHP.eB. (**C**) Vector genome (vg) biodistribution of AAV-PHP.eB or AAV9 two hours after intravascular administration to C57BL/6J or BALB/cJ mice (n=6/virus/line, mean±s.e.m.; 2-way ANOVA; *p<0.05, **p<0.01, ***p<0.001). (**D**) *Ly6a* and *Ly6c1* SNPs correlate with the nonpermissive phenotype. Missense SNPs relative to C57BL/6J are listed as the amino acid change. SRV, splice region variant; IV, intron variant; SDV, splice donor variant. (**E**) Expression data (mean fragments per kilobase-million ± s.d.) for *Ly6a, Ly6c1*, and *Pecam1*^*23*^.

In light of this disparity, we sought to test for the enhanced CNS tropism of the AAV-PHP.B capsids across a panel of mouse lines, harnessing the natural genetic variation between mice to identify the genetic variants and, subsequently, candidate gene(s) responsible for the difference in CNS transduction by AAV-PHP. eB. Diverging from conventional diversity outbred or genetic linkage studies requiring the inter-breeding of mouse lines, we leveraged the open-source software Hail^21^ to perform a genome-wide association study (GWAS) utilizing the Mouse Genome Project database spanning 36 mouse lines^22^. We aimed to rapidly identify variants whose alleles segregate across mice with the observed phenotype (permissive or non-permissive to enhanced CNS transduction by AAV-PHP.eB). Starting from millions of genetic variants comprised of single-nucleotide polymorphisms (SNPs) as well as insertions and deletions (indels), we narrowed our analysis to variants predicted to affect expression, splicing, or protein coding regions (Supplementary Table 1). Using a statistical simulation framework, we estimated that 12 mouse lines would be sufficient to narrow our search to ∼10 high/medium impact variants (Supplementary Fig. 2, Supplementary Table 1), which could feasibly be experimentally interrogated for the enhanced AAV-PHP.eB CNS tropism. Based on this estimation, we acquired mice from 13 commercially available lines, including C57BL/6J and BALB/cJ, and administered 10^11^ vector genomes (vg)/animal of AAV-PHP.eB, which packaged an AAV genome encoding a green fluorescent protein (GFP) with a nuclear localization signal (NLS-GFP). Intravenous administration of AAV-PHP.eB resulted in GFP expression in the brain of permissive lines such as C57BL/6J, but not those of nonpermissive mice such as BALB/cJ; we identified seven permissive and six nonpermissive lines (Supplementary Fig. 2).

The GWAS reduced the number of high/medium impact gene variants to missense SNPs in the *Ly6a* and *Ly6c1* genes (Fig. 1D). RNA sequencing data from sorted mouse brain cells (www.BrainRNAseq.org)^23^ indicates that *Ly6a* and *Ly6c1* are expressed in brain endothelial cells (Fig. 1E). Intriguingly, the mouse Ly6 locus is linked to susceptibility to mouse adenovirus (MAV1)^24^, which possesses an endothelial cell tropism that causes fatal hemorrhagic encephalomyelitis in C57BL/6 but not BALB/ cJ mice^25^. The *Ly6* gene family also influences susceptibility to infection by HIV1^26,27^, Flaviviridae (yellow fever virus, dengue, and West Nile virus^28^, Influenza A^29^, and Marek’s disease virus in chickens^30^. Despite these promising links between *Ly6* genes and various virus susceptibilities, the mechanism remains elusive and has not been leveraged in the development of gene therapy vectors.

We first validated whether protein expression or localization differences arising from genetic variation within *Ly6a* or *Ly6c1* is associated with the differential AAV-PHP.eB tropism across mouse lines. Immunohistochemistry (IHC) assays revealed that LY6A is abundant within the CNS endothelium of C57Bl/6J mice but not in BALB/cJ mice (Fig. 2A-C). In contrast, *Ly6c1* was expressed on the CNS endothelium of both lines (Fig. 2D). The reduced level of LY6A, relative to that observed in C57BL/6J mice, correlated with the nonpermissive AAV-PHP.eB transduction phenotype across five of six mouse strains (Supplementary Fig. 3). This suggested that reduced LY6A levels may contribute to, but is unlikely to be the only factor responsible for the absence of AAV-PHP.eB transduction. LY6A was evident as multiple bands by western blotting, but only the more slowly migrating band was detectable at low levels in BALB/cJ mice (Fig. 2E), suggesting that LY6A maturation or post-translational processing differs between the mouse lines. We noted that the Ly6a V106A, *Ly6c1* P133L (Fig. 1D), and several noncoding SNPs in the *Ly6* locus perfectly segregate with the nonpermissive phenotype (observed in six out of six nonpermissive mouse strains). Together, these results suggest an association between Ly6 gene variants and permissivity to transduction by AAV-PHP.eB.

**Figure 2.**
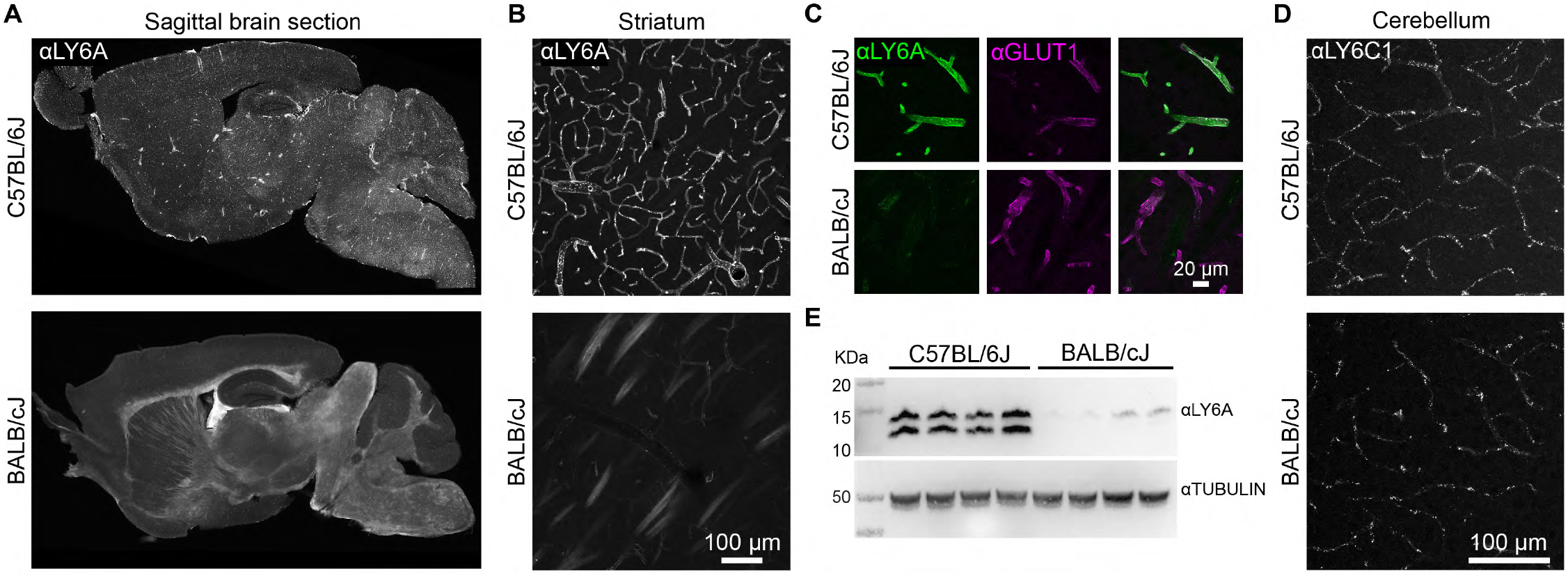
*Ly6a* expression is altered in the BALB/cJ mouse brain. (**A**) Representative images of LY6A IHC in sagittal brain sections from C57BL/6J or BALB/cJ mice. LY6A IHC in the striatum (**B**, white or **C**, green) with IHC for GLUT1 (magenta) to mark endothelial cells (see Supplementary Fig. 3 for images of LY6A IHC in brain sections from the 11 additional mouse lines). LY6C1 IHC in the cerebral cortex (**D**) of C57BL/6J or BALB/cJ mice. (**E**) Western blot from forebrain lysates reveals LY6A abundance and protein states in each mouse line.

### *Ly6a* is necessary for the enhanced CNS transduction phenotype of AAV-PHP.eB

We sought to determine whether LY6A and/or LY6C1 are necessary for the ability of AAV-PHP.eB to bind and transduce CNS endothelial cells. To achieve this, we performed *Ly6a* and *Ly6c1* knockout experiments in brain microvascular endothelial cells (BMVECs) from C57BL/6J mice, which express both genes (Fig. 3A) and are more efficiently bound and transduced by AAV-PHP.eB than by AAV9 (Fig. 3B, C). We used CRISPR/ SaCAS9^31^ and *Ly6a-* or *Ly6c1*-specific sgRNAs to disrupt each gene. Using three different sgRNAs to target *Ly6a*, we achieved a ∼50% reduction of LY6A (Supplementary Fig. 3) and a 50% reduction in binding by AAV-PHP.eB, but not AAV9 (Fig. 3D and Supplementary Fig. 4). A similar reduction in transduction by AAV-PHP.eB was observed (Fig. 3E); AAV9 transduction of BMVECs was inefficient and not included for comparison. None of the sgRNAs targeting *Ly6c1* affected AAV-PHP.eB or AAV9 binding to the BMVECs (Fig. 3D). Collectively, the reduced AAV-PHP.eB binding resulting from *Ly6a* disruption in BMVECs, the high level of *Ly6a* expression within the CNS endothelium of permissive mouse lines, and the association of *Ly6a* SNPs with the nonpermissive phenotype, suggest that LY6A functions as a receptor for AAV-PHP.eB.

**Figure 3.**
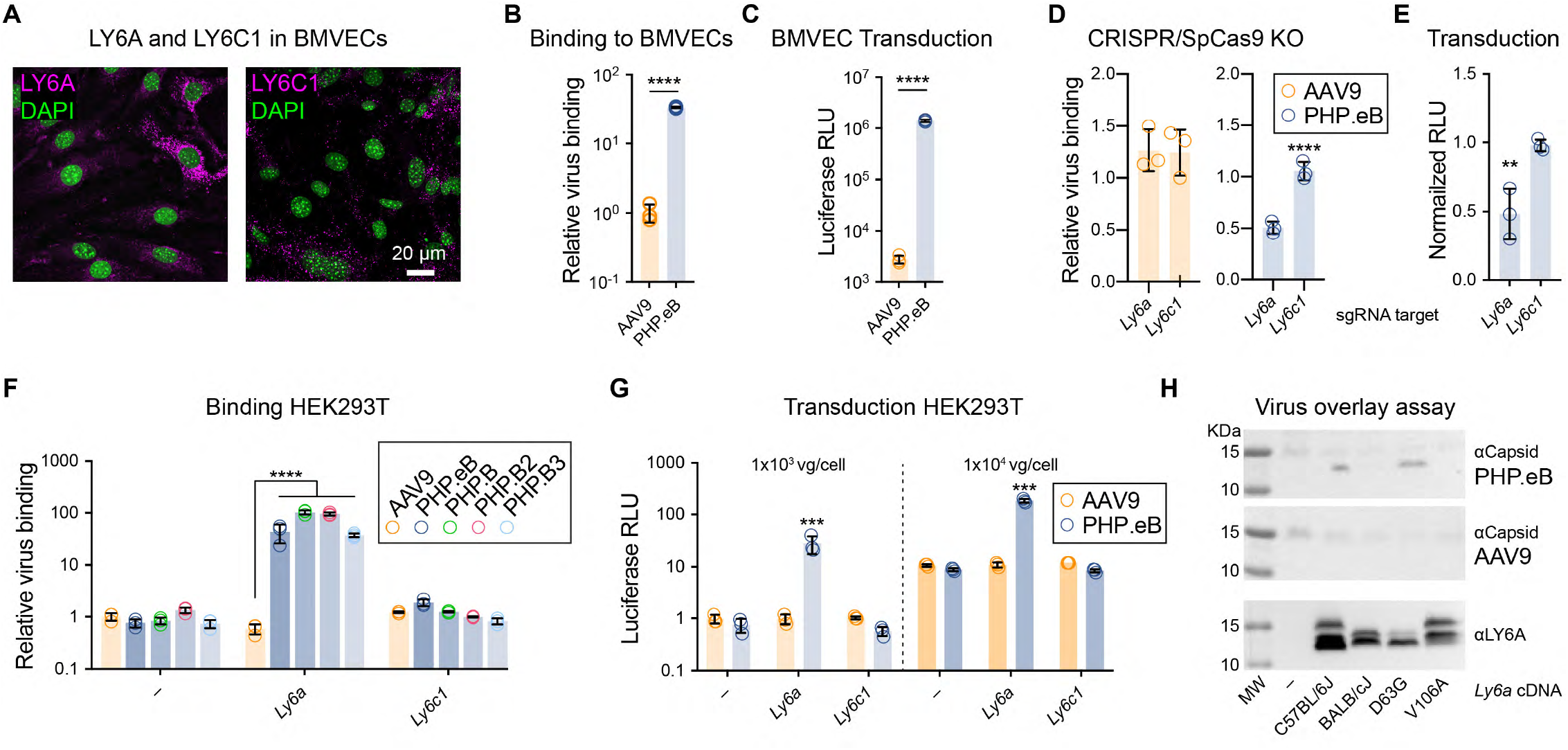
*Ly6a* is necessary and sufficient for enhanced binding and transduction by AAV-PHP.eB. (**A**) LY6A (magenta, left) and LY6C1 (magenta, right) immunostaining with nuclei (green, DAPI) in BMVECs. AAV9 and AAV-PHP.eB binding (**B**) and transduction (**C**) of BMVECs (n=3/virus, mean±S.D., ****p<0.0001; two-tailed t test). Binding was assessed by qPCR of the viral genome. Transduction was assessed by measuring Luciferase luminescence in relative light units (RLU). Binding (**D**; 2-way ANOVA, Dunnett’s multiple comparison test) and transduction (**E**; 1-way ANOVA, Sidak’s post test) by the indicated virus in cells treated with a vector containing an sgRNA to disrupt *Ly6a* or *Ly6c1* or no sgRNA (n=3/group, mean±S.D. **p<0.01, ****p<0.0001). Each data point represents cells that received a different sgRNA. (**F**) Binding of the indicated virus to HEK293T cells transfected with *Ly6a, Ly6c1*, or mock (–). (**G**) Transduction measured by Luciferase assay normalized to AAV9 on mock transfected cells. (**F**, **G**; n=3/group, **p<0.01, ****p<0.0001; 2-way ANOVA, Tukey’s correction). (**H**) Virus overlay assay using lysates from HEK293T cells transfected with *Ly6a* cDNAs from C57BL/6J or containing one or both BALB/cJ SNPs. Panels show immunoblotting for AAV capsid proteins after overlaying with AAV-PHP.eB or AAV9. Bottom panel shows the same blot probed with αLY6A.

### *Ly6a* expression enhances transduction by AAV-PHP.eB in human cells

We next asked whether ectopic *Ly6a* expression is sufficient for increased binding and transduction by AAV-PHP.eB. We transiently transfected HEK293T cells with cDNAs encoding C57BL/6J *Ly6a* or *Ly6c1*, and evaluated the effects on binding and transduction by AAV-PHP.B capsids. Remarkably, *Ly6a* expression resulted in a >50-fold increase in binding by each of the AAV-PHP.B capsids to HEK293T cells, but did not increase binding by AAV9 (Fig. 3F). As expected, expression of *Ly6a*, but not *Ly6c1*, enhanced transduction by AAV-PHP.eB by 30-fold compared to the untransfected control (Fig. 3G).

### Binding between LY6A and AAV-PHP.eB is disrupted by a V106A variant of LY6A

To determine whether AAV-PHP.eB directly binds LY6A and whether either of the missense SNPs (D63G or V106A) in the BALB/cJ *Ly6a* gene (Fig. 1D) affect binding to AAV-PHP.eB, we performed virus overlay assays^32^. AAV-PHP.eB bound a protein that co-migrates with LY6A (Fig. 3H) from HEK293T cells transfected with the C57BL/6J or D63G *Ly6a* cDNAs, but not from cells expressing *Ly6a* from the BALB/cJ or V106A cDNAs. Interestingly, the V106A variant is located near the predicted cleavage and GPI anchoring site (ω), and is predicted to reduce the likelihood of GPI-anchor modification^33^ (Supplementary Table 2).

### LY6A enhances AAV-PHP.eB transduction independently of known AAV9 receptors

To determine whether LY6A acts solely as a primary attachment factor or has additional roles in promoting the internalization and trafficking of AAV-PHP.eB, we explored whether AAV-PHP. eB binding and transduction are dependent on known receptor interactions. AAVs typically use multiple cellular receptors for attachment and internalization and intracellular trafficking^34^; AAV9 utilizes galactose as an attachment factor^35^, and, like most AAVs, requires the AAV receptor (AAVR) for intracellular trafficking and transduction^32^.

We first tested the effect of varying levels of galactose on cell surface glycoproteins on LY6A-mediated AAV-PHP.eB binding. Using Pro5 Chinese Hamster ovary (CHO) cells and derivatives that expose excess galactose (Lec2 cells) or are unable to add galactose to its glycoproteins (Lec8 cells)^36^, which were previously used to map the galactose binding site on the AAV9 capsid^35^, we assessed whether AAV-PHP.eB-LY6A interactions are influenced by galactose levels. As expected, AAV9 and AAV-PHP.B similarly bound and transduced Lec2 cells more efficiently than Lec8 or Pro5 cells (Fig. 4A-C), showing that AAV-PHP.B also utilizes galactose for cell attachment. In contrast, ectopic *Ly6a* expression selectively increased AAV-PHP.eB, but not AAV9, binding and transduction (Fig. 4A, B) to Pro5 and Lec8 cells. *Ly6a* expression did not increase binding of AAV-PHP.eB to Lec2 cells (Fig. 4A), potentially due to the high levels of binding driven by excess galactose. Interestingly, *Ly6a* expression enhanced AAV-PHP.eB transduction of Pro5, Lec2, and Lec8 cells (Fig. 4B). The finding that *Ly6a* expression rendered Lec8 cells as receptive to AAV-PHP. eB transduction as Pro5 cells indicates that LY6A functions as an attachment factor for AAV-PHP.eB independently of galactose. Furthermore, *Ly6a* expression enhanced AAV-PHP.eB transduction of Lec2 cells without increasing binding, suggesting that LY6A mediates the internalization and/or trafficking of AAV-PHP.eB.

**Figure 4.**
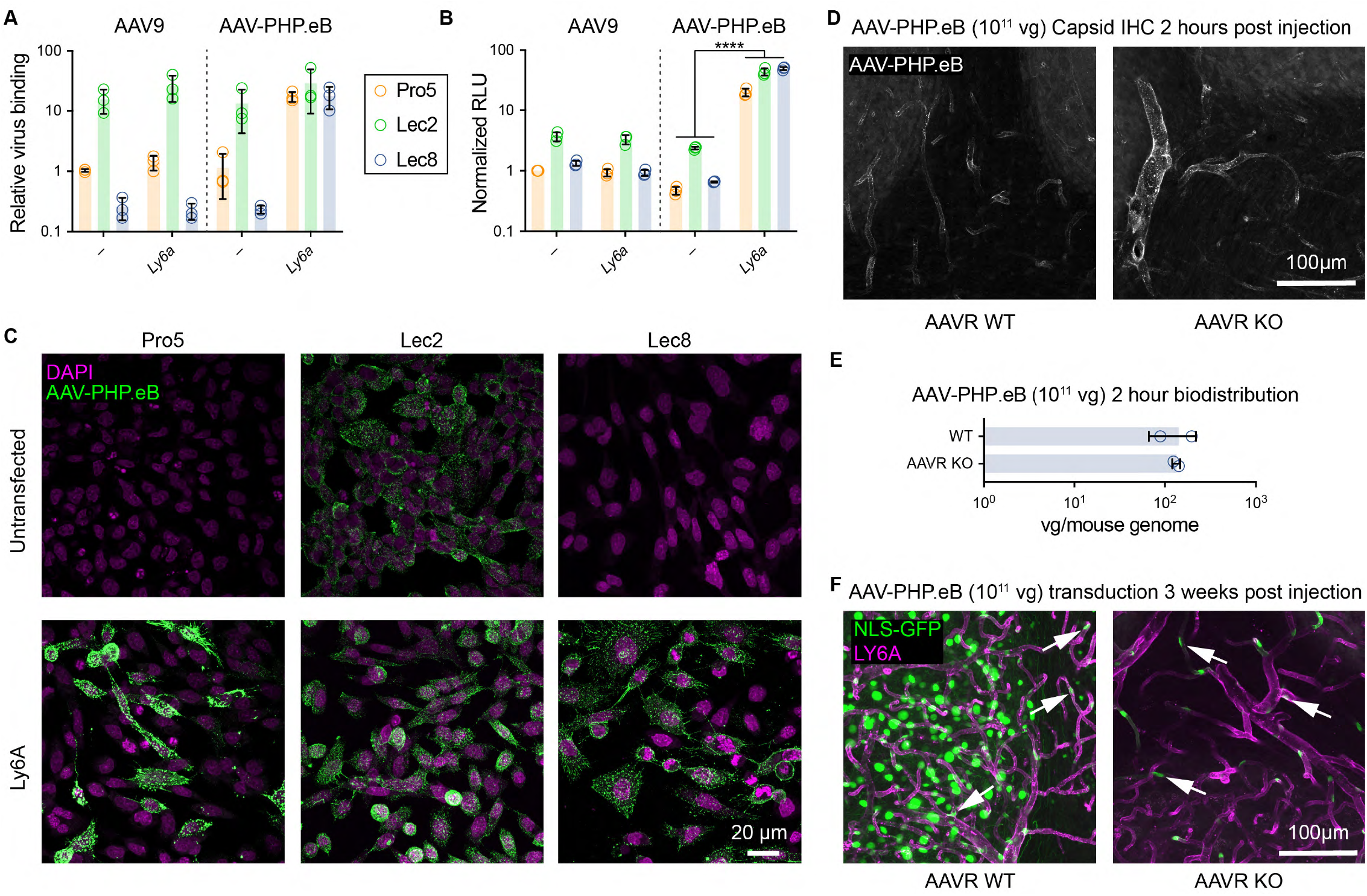
*Ly6a* expression renders cells permissive to AAV-PHP.eB transduction in the absence of galactose or AAVR. (**A-C**) AAV-PHP.eB or AAV9 viruses were added to control Pro5 CHO cells, Lec2 CHO cells with excess galactose, or Lec8 CHO cells deficient for galactose transfer. (**A**) Quantification of AAV binding to CHO cell derivatives via qPCR for viral genomes. (**B**) Transduction of CHO cells as measured by Luciferase assay 48 hours after virus addition, normalized to values from Pro5 cells transduced with AAV9. (**C**) AAV-PHP. eB capsid immunostaining (magenta) of CHO cells that were untransfected (top row) or transfected with *Ly6a* (bottom row). (**D**-**F**) AAVR WT or KO mice were intravenously injected with AAV-PHP.eB:CAG-*NLS-GFP* (10^11^vg/mouse) and brain tissue was assessed via IHC for capsid binding (**D**) or vector genome biodistribution analysis (**E**) at two hours or transduction (**F**) at three weeks post injection. (**D**-**F**) (n=2 per group/per experiment, mean ± SD). (**F**) Representative images of GFP fluorescence (green) with LY6A IHC (magenta).

In this vein, we reasoned that this interaction may render *Ly6a*-expressing cells permissive to AAV-PHP.eB transduction in the absence of AAVR, which is essential for the intracellular trafficking of nearly all AAV capsids including AAV9^37^. To test this possibility, AAVR WT and KO FVB/NJ mice^37^ were injected with AAV-PHP.eB, and their brains were collected two hours later for capsid detection. We detected AAV-PHP.eB along the vasculature and in the brain lysates of both AAVR WT and KO mice (Fig. 4D, E). Three weeks post-administration, AAV-PHP.eB transduction of neurons and astrocytes, which do not express *Ly6a*, was nearly absent in the brains of AAVR KO mice. In contrast, AAV-PHP. eB transduced *Ly6a*-expressing endothelial cells throughout the brains of both AAVR WT and KO mice (Fig. 4F). Collectively, these results demonstrate that the AAV-PHP.eB virus uses a unique and unexpected endothelial cell receptor, LY6A, to gain access to the CNS.

## Discussion

Our development of AAV-PHP.B capsids with dramatically improved tropism for the mouse CNS provided a proof-of-concept that AAVs can be engineered to cross the BBB^5,6^. In this publication, we leveraged Hail^21^, the Mouse Genome Project dataset^22^, and 13 commercially available mouse lines to rapidly identify a single amino acid change, out of a starting pool of millions of genetic variants, that impacts AAV-PHP.eB permissivity; the code was implemented and run end-to-end on whole-genome sequencing data within hours, harnessing Hail’s ability to scale computation across a large compute cluster, and the *in vivo* screening was completed in three weeks. The speed and reduced number of animals required for this approach is unprecedented compared to the conventional approaches of using diversity outbred lines or breeding generations of mice to determine the approximate genomic loci that segregates with a given phenotype.

Our identification of LY6A as the receptor engaged by the AAV-PHP.B capsids has important translational value. We demonstrate that LY6A facilitates both binding and internal trafficking of AAV-PHP.B capsids. Other cellular factors that share LY6A’s properties such as abundant luminal surface exposure on brain endothelium, localization within specific lipid micro-domains through GPI anchoring, or specific recycling/intracellular trafficking capabilities, may be prime molecular targets for gene delivery vectors in mice, NHPs, and humans. Although *Ly6a* is absent in primates, other LY6 proteins with homologs in primates are present within the CNS endothelium, and can potentially be explored and harnessed for AAV capsid engineering. Directing capsids and/or other biologicals that target these receptors can open up new therapeutic avenues for treating a wide range of currently intractable neurological diseases.

## Acknowledgements

We thank R. Hosking, T. Brown, and F.-E. Eid for helpful comments on the manuscript; E. Dzhura and T. Bates for lab organization; G. Fishell and N. Yusuf for use of the Zeiss microscopes. This study was supported by the Stanley Foundation and Stanley Center for Psychiatric Research, and an NIH Somatic Cell Genome Editing Consortium grant (NINDS UG3 NS111689-01).

## Author contributions

B.D. and Q.H. designed and implemented the study; B.D. wrote the paper with significant input from Y.A.C.; Q.H., K.C., I.T., B.D., and C.B. performed experiments; Q.H., B.D., T.P., J.B., and C.S. analyzed data; R.D. and A.B. helped to plan experiments and provided data and reagents. All authors discussed the results and contributed to the manuscript.

## Supplementary Materials

Materials and Methods

Supplementary Figures 1 – 4

Supplementary Tables 1 – 3

## Materials and Methods

### Mouse strain permutation analysis

Access to whole-genome sequencing data for 36 commercially available mouse lines^22^ made it possible to estimate the number of lines necessary to produce a shortlist of candidate variants. Starting from the known permissive C57BL/6J and nonpermissive BALB/cJ strains, we used the genomic analysis software Hail^21^ to simulate permissivity phenotypes for other mouse lines and compute the number of candidate variants in both the high and high-and-medium predicted functional impact classes. First, we sampled the probability p that a random mouse line is permissive from a Beta(α =2, β=2) distribution, based on the two known mouse phenotypes. Second, we sampled a subset of additional commercially available lines (Supplementary Table 3). Third, we simulated a permissivity phenotype for each line in the subset by flipping a p-coin. Finally, we calculated the number of variants with perfect allelic segregation between the permissive and nonpermissive lines. We ran 500 iterations of this model for each of ten subset sizes (3 to 12 mice), providing a distribution over the number of candidate variants at each mouse sample size. This simulation informed our decision regarding the number of mouse lines to order and test in parallel.

### Plasmids and primers

The AAV-PHP.eB Rep-Cap trans plasmid was generated by gene synthesis (GenScript). AAV9, AAV-PHP.B, AAV-PHP.B2, and AAV-PHP.B3 were generated by replacing the AAV-PHP.eB variant region with that of AAV9, AAV-PHP.B, B2, or B3 using isothermal HiFi DNA Assembly (NEB). The reporter and Ly6 expression vectors were cloned into an AAV-CAG-WPRE-hGH pA backbone obtained from Viviana Gradinaru through Addgene (#99122). GFP, 2A-luciferase, *Ly6a* and *Ly6c1* (splice variant 1) cDNAs were synthesized as gBlocks (IDT). The NLS-GFP were synthesized using the N-terminal SV40 NLS sequence present in the Addgene plasmid #99130. The CMV-SaCAS9 vector (AAV-CMV∷NLS-SaCas9-NLS-3xHA-bGHpA;U6∷BsaI-sgRNA) was obtained from Dr. Feng Zhang through Addgene (#61591). sgRNAs specifically targeting *Ly6a* or *Ly6c1* were cloned after the U6 promoter using a single bridge oligo for each reaction as recommended (HiFi DNA Assembly, NEB). The Broad GPP sgRNA tool for SaCAS9 was used to identify suitable SaCAS9 target sites^38^.

The following primers were used for *Ly6a* sgRNA cloning: 5’-CTTGTGGAAAGGACGAAACACCGAATTACCT-GCCCCTACCCTGAGTTTTAGTACTCTGGAAACAG, 5’-CTTGTGGAAAGGACGAAACACCGCTTTCAATATTAG-GAGGGCAGGTTTTAGTACTCTGGAAACAG, 5’-CTTGT-GGAAAGGACGAAACACCGAATATTGAAAGTATGGAGATC-GTTTTAGTACTCTGGAAACAG

The following primers were used for *Ly6a* sgRNA cloning: 5’-CTTGTGGAAAGGACGAAACACCGACTG-CAGTGCTACGAGTGCTAGTTTTAGTACTCTGGAAACAG, 5’-CTTGTGGAAAGGACGAAACACCGCAGTTACCTGCCGCG-CCTCTGGTTTTAGTACTCTGGAAACAG, 5’-CTTGTGGAAAG-GACGAAACACCGGATTCTGCATTGCTCAAAACAGTTTTAG-TACTCTGGAAACAG

The following primers were used for biodistribution assays and/ or binding assays

GFP: 5’-TACCCCGACCACATGAAGCAG, 5’-CTTGTAGTTGCCGTCGTCCTTG

Mouse *Gcg*: 5’-AAGGGACCTTTACCAGTGATGTG, 5’-ACTTACTCTCGCCTTCCTCGG

Human *GCG*: 5’-ATGCTGAAGGGACCTTTACCAG, 5’-ACTTACTCTCGCCTTCCTCGG

CHO *Gcg*: 5’-ATGCTGAAGGGACCTTTACCAG, 5’-CTCGCCTTCCTCTGCCTTT

### CRISPR/SaCas9 KO experiments

AAV-PHP.eB vectors with sgRNA sequences targeting *Ly6a* and *Ly6c1* were generated and purified to knockout the respective genes in C57BL/6 mouse primary brain microvascular endothelial cells (CellBiologics, Cat. #C57-6023). AAV vectors (1×10^6^vg/cell) were used to transduce cells every 3 days for 3 rounds to achieve higher knockout efficiency. Cells were passaged as necessary.

### Cell lines and primary cultures

HEK293T/17 (CRL-11268), Pro5 (CRL-1781), Lec2 (CRL-1736), and Lec8 (CRL-1737) were obtained from ATCC. BMVEC cells were obtained from Cell Biologics (C57-6023) and cultured as directed by the manufacturer.

### Virus production and purification

Recombinant AAVs were generated by triple transfection of HEK293T cells (ATCC CRL-11268) using polyethylenimine (PEI) and purified by ultracentrifugation over iodixanol gradients as previously described^6^.

### Western blotting and virus overlay assays

The virus overlay assay was performed as previously reported^32^ with some modifications. Briefly, protein lysates were separated on Bolt 4-12% Bis-Tris Plus gels and transferred onto nitrocellulose membranes. After incubation with AAV9 or PHP.eB at 5e11 vg/ mL, the membranes were fixed with 4% PFA at room temperature for 20 minutes to crosslink the interaction between the capsid and its target protein, followed by 2M HCl treatment at 37°C for 7 minutes to expose the internal capsid epitope for detection. The blots were then rinsed and incubated with anti-AAV VP1/VP2/VP3 (1:20; American research products, Inc., cat. #03-65158) followed by incubation with a horseradish peroxidase (HRP)-conjugated secondary antibody at 1:5000. The detection of the HRP signal was by SuperSignal West Femo Maximum Sensitivity Substrate using a Bio-Rad ChemiDoc ™ MP system #1708280.

### Animals

All procedures were performed as approved by the Broad Institute IACUC or Massachusetts General Hospital IACUC (AAVR experiments). AKR/J (000648), BALB/cJ (000651), CBA/J (000656), CAST/EiJ (000928), C57Bl/6J (000664), C57BL/J (000668), DBA/2J (000671), FVB/NJ (001800), LP/J (000676), MOLF/EiJ (000550), NOD/ShiLtJ (001976), NZB/ B1NJ (000684), and PWK/PhJ (003715) were obtained from the Jackson Laboratory (JAX). AAVR mice were a generous gift from Dr. J.E. Carette (Stanford) to Dr. Balazs and have been previously described^37^. Recombinant AAV vectors were administered intravenously via the retro-orbital sinus in young adult male or female mice. Mice were randomly assigned to groups based on predetermined sample sizes. No mice were excluded from the analyses. Experimenters were not blinded to sample groups.

### Tissue processing, immunohistochemistry and imaging

Mice were anesthetized with Euthasol (Broad) or ketamine (MGH) and transcardially perfused with phosphate buffered saline (PBS) at room temperature followed by 4% paraformaldehyde (PFA) in PBS. Tissues were post-fixed overnight in 4% PFA in PBS and sectioned by vibratome. IHC was performed on floating sections with antibodies diluted in PBS containing 10% donkey serum, 0.1% Triton X-100, and 0.05% sodium azide. Primary antibodies were incubated at room temperature overnight. The sections were then washed and stained with secondary (Alexa-conjugated antibodies, 1:1000) for 4 hours or overnight. Primary antibodies used were mouse anti-AAV capsid (1:20; American Research Products, 03-65158, clone B1), LY6A (1:250; BD Bioscience, 553333 or ThermoFisher, 701919), LY6C1 (1:250; Millipore-Sigma, MABN668), Glut1 (1:250; Millipore Sigma, 07-1401). To expose the internal B1 capsid epitope in intact capsids, tissue sections or cells on coverslips were treated for 15 or 7 minutes, respectively, with 2M HCl at 37°C. The treated samples were then washed extensively with PBS prior to addition of the primary antibody.

### *In vivo* vector and capsid biodistribution

5- to 6-week-old C57Bl/6J mice, BALB/cJ mice, AAVR WT FVB/ NJ, or AAVR KO FVB/NJ mice were injected intravenously with 10^11^vg of AAV vector packaged into the indicated capsid. One or two hours after injection, the mice were perfused with PBS and tissues were collected and frozen at −80°C. Samples were processed for AAV genome biodistribution analysis by measuring the number of vector genomes (vg) in the sample using absolute qPCR for the GFP sequence. The values are reported as vg/mouse genome after normalizing to the number of copies of host cell genomes using primers recognizing mouse *Gcg* or human *GCG* as previously described^6^. To visualize the capsid distribution, mice were perfused with 4% PFA after dosing with AAV vector and brains were sectioned into 100 µM and labeled with the indicated antibodies.

### Microscopy

Images were taken on an Axio Imager.Z2 Basis Zeiss 880 laser scanning confocal microscope fitted with the following objectives: PApo 10x/0.45 M27, Plan-Apochromat 20x/0.8 M27, or Plan-APO 40x/1.4 oil DIC (UV) VIS-IR or on a Zeiss Imager.M2 (5x tiled, whole sagittal section images). All images that were compared within an experiment were acquired and processed under identical conditions.

### *In vitro* binding assays

*Ly6* family members (0.5 µg/well) were transfected into HEK293T cells (3×10^5^/well) using PEI or into CHO cells (1.5×10^5^/well) with lipofectamine 3000 reagent (ThermoFisher, L3000001) in 24-well plates. 48 hours later, the cells were chilled to 4°C and the media was exchanged with fresh cold media containing the indicated recombinant AAV (10^5^ vg/cell). One hour later, cells were washed three times with cold PBS, then fixed with 4% PFA for IHC or lysed for genomic DNA extraction and qPCR analyses. Vector genomes bound were normalized to the number of cell genomes by using primers specific to *Gcg or GCG*. For BMVECs, 2×10^4^ cells/well were seeded in 12-well plates the day before exposure to virus. The assay was performed as above except that the AAV vectors were added at 10^6^vg/cell.

### Luciferase transduction assay

*Ly6* family members (0.1 µg/well) were transfected into the indicated cells (HEK293/17: 4×10^5^/well; CHO: 2.5×10^4^/well, BMVECs: 5×10^3^/well) in 96-well plates (PerkinElmer, 6005680) in triplicate. 48 hours later, cells were transduced with AAV-CAG-GFP-2A-Luciferase-WPRE packaged into AAV9 or AAV-PHP.eB. Luciferase assays were performed with Britelite plus Reporter Gene Assay System (PerkinElmer, 6066766). Luciferase activity was reported as relative light units (RLU) as raw data or normalized to non-transfected control wells transduced with AAV9, or a control transduced without a sgRNA (Fig. 3E).

### Statistical analysis and experimental design

Microsoft Excel and Prism 8 were used for data analysis. For the comparison between AAV9 and AAV-PHP.eB biodistribution, a group size of 6 per group (3 males and 3 females) was used based on prior data that indicated a large effect size. No animals or samples were excluded from the analysis. Supplementary Fig. 1 images are representative of two animals per group. To evaluate AAV-PHP.eB in the 13 mouse lines, AAV9 (n=1, 10^11^vg/animal) or AAV-PHP.eB (n=2; one per dose at 10^11^ and 10^12^vg/animal). LY6A IHC in Supplementary Fig. 3 is representative of two animals/line. *In vitro* transduction and binding experiments display the means from three independent experiments. In Fig. 3D and Fig. 3E, each data point represents a different sgRNA, each averaged from three independent experiments. Data were normalized separately for each AAV capsid to cells transduced with SpCas9 vectors without an sgRNA. Supplementary Fig. 4 presents the same data as Fig. 3D separated by each individual sgRNA. Data from AAVR WT and KO mice are representative of two mice per genotype per time point post-injection. In all panels, *p<0.05, **p<0.01, ***p<0.001, ****p<0.0001.

---

**Supplementary Figure 1.**
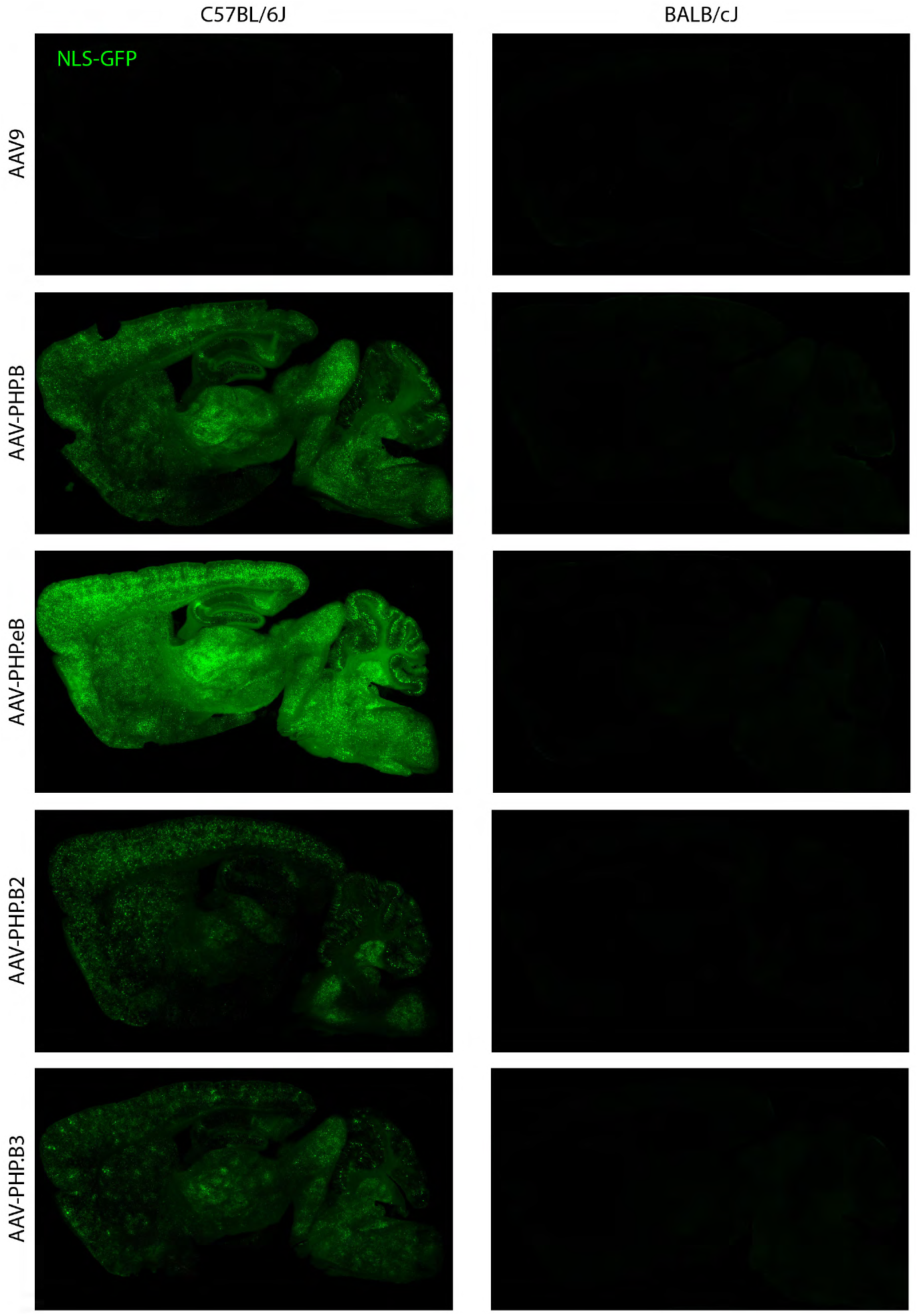
Transduction of the brain of BALB/cJ and C57BL/6J mice by AAV9, AAV-PHP.B, AAV-PHP.eB, AAV-PHP.B2, or AAV-PHP.B3. Images of GFP fluorescence in whole brain sagittal sections from C57BL/6J (left column) or BALB/cJ (right column) two weeks after intravenous injection of 10^11^vg/mouse AAV-CAG-NLS-GFP packaged into the indicated capsid.

**Supplementary Figure 2.**
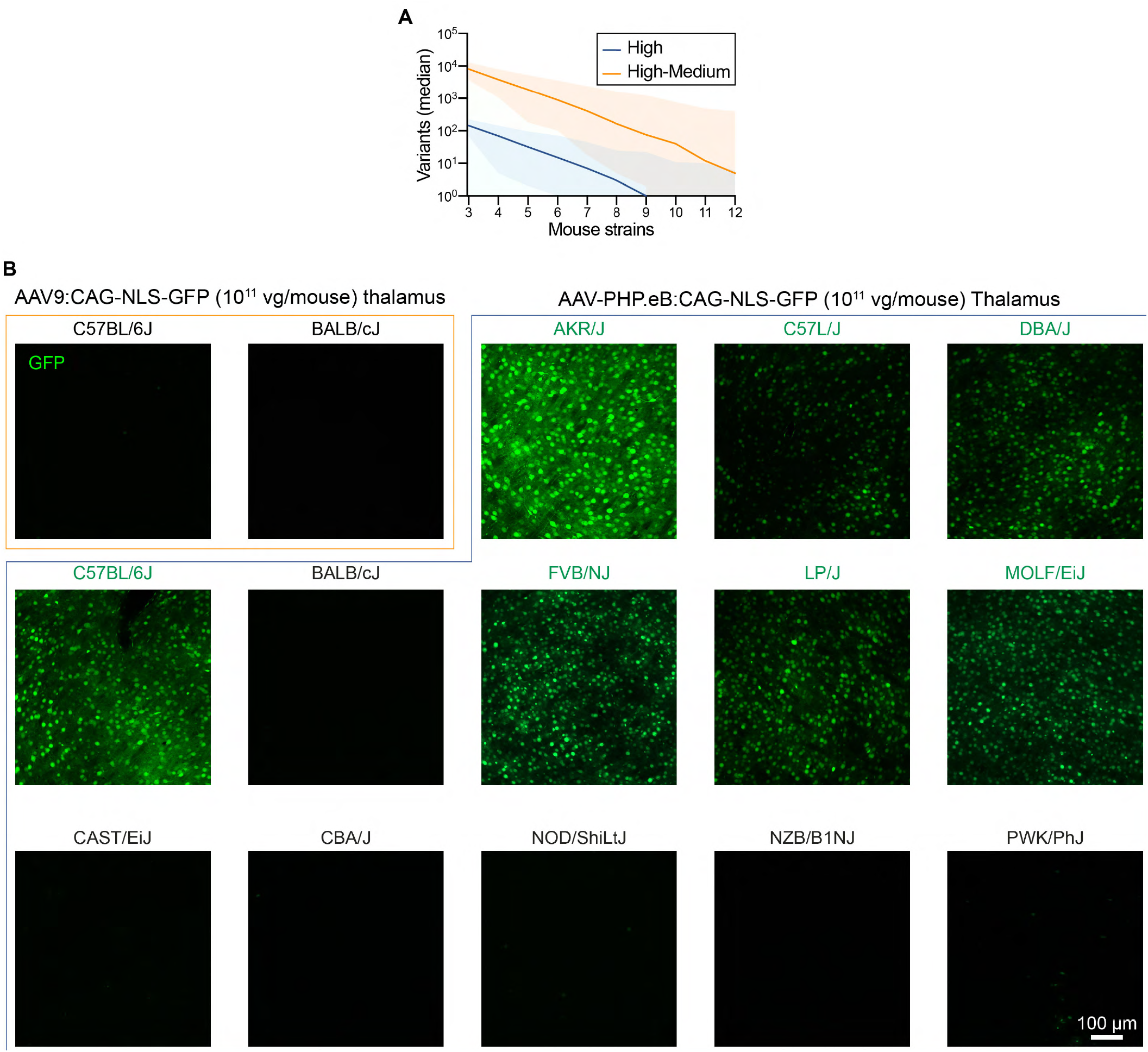
The AAV-PHP.eB CNS tropism phenotype is absent in a subset of mouse lines. (**A**) The number of mouse lines required to reduce the number of candidate variants associated with AAV-PHP. eB permissivity. The plotted lines depict the median number of simulated candidate variants; high (loss-of-function; blue) or high+medium (loss-of-function, missense, splicing variant; orange). Shaded regions represent 5–95th percentiles. (**B**) Native GFP fluorescence in the mouse thalamus two weeks after intravenous injection of 10^11^vg/mouse CAG-NLS-GFP packaged into AAV9 (first two panels from top left) or AAV-PHP.eB.

**Supplementary Figure 3.**
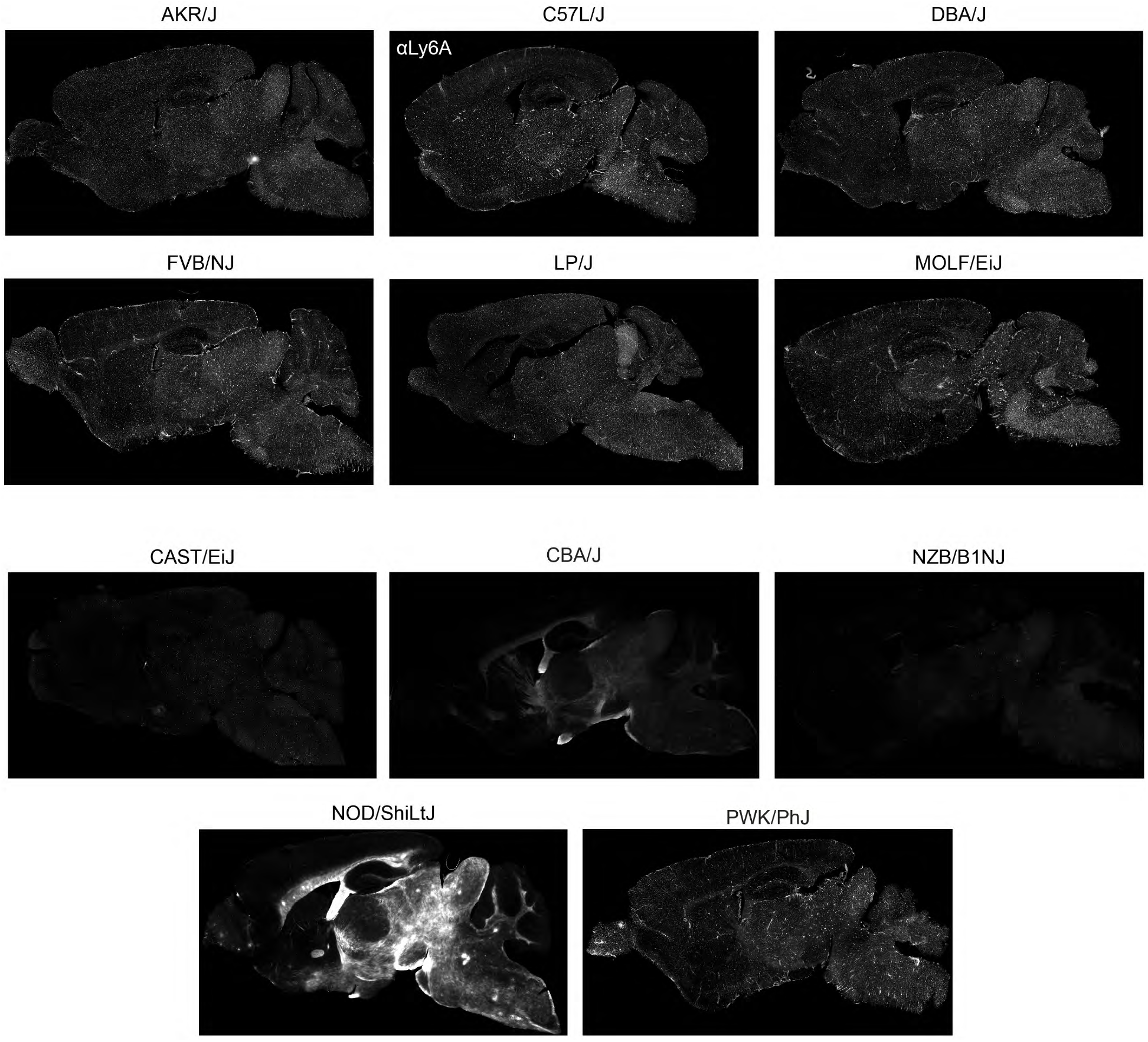
LY6A is highly abundant on the brain endothelium of permissive mouse lines, but its distribution and/or expression is altered in mice nonpermissive to AAV-PHP.eB CNS transduction. Sagittal whole brain images show LY6A IHC in several representative permissive and nonpermissive mouse lines. LY6A was localized in white matter tracts within the CNS in a subset of the nonpermissive mouse lines (BALB/cJ, CBA/J and NOD/ShiLtJ) as has been previously reported^39,40^.

**Supplementary Figure 4.**
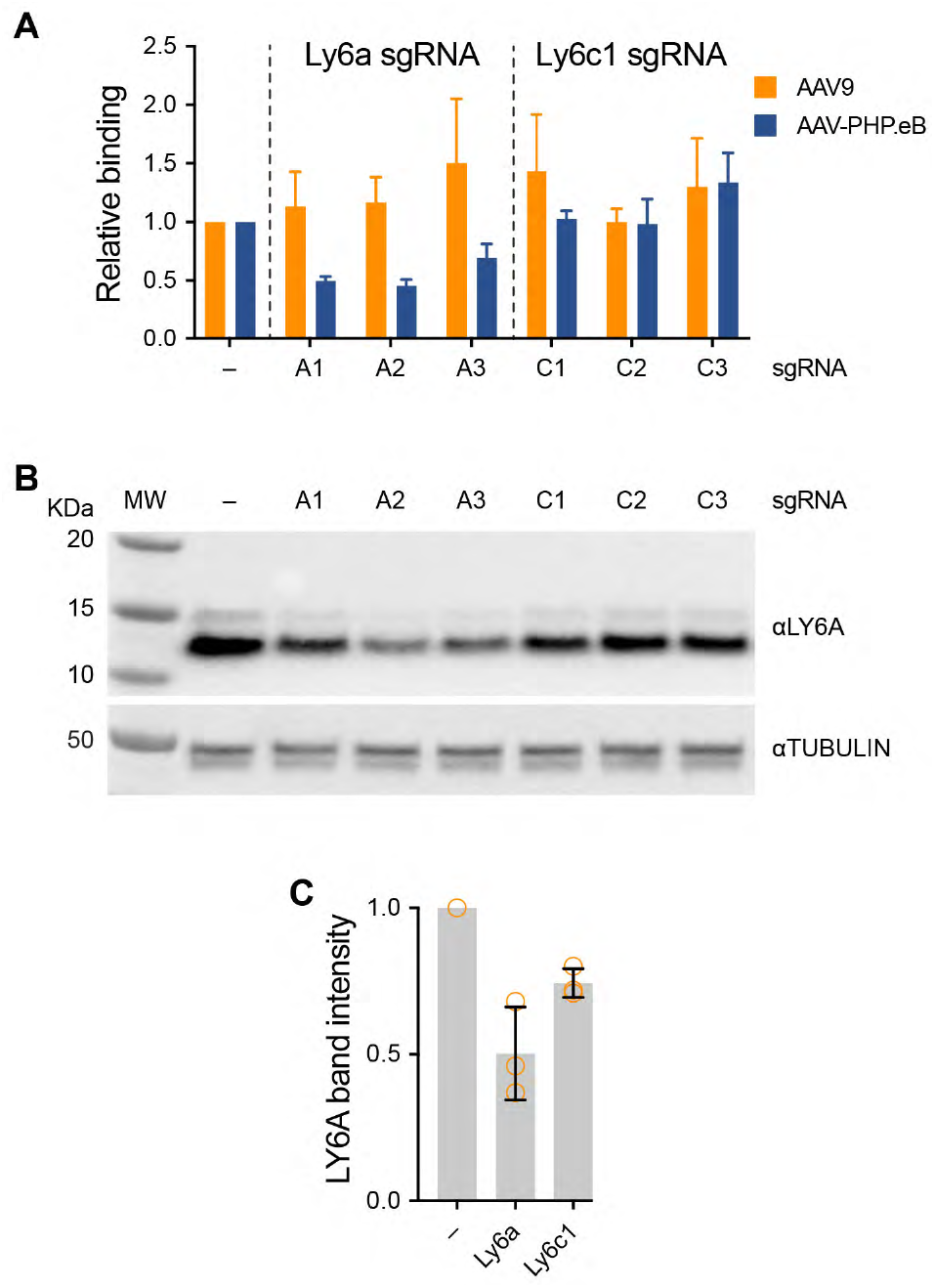
Disruption of *Ly6a* and *Ly6c1* using CRISPR/Cas9 and target-specific sgRNAs results in reduced LY6A protein and reduced AAV-PHP.eB binding. (**A**) The individual sgRNA data used to generate Fig. 3D is plotted. AAV9 and AAV-PHP.eB binding data were normalized individually to control samples from cells treated with a control SaCAS9 vector lacking a sgRNA (–). (**B**) Western blots for LY6A (top) or TUBULIN (bottom) in lysates prepared from BMVECs treated with the individual sgRNAs shown in (**A**). (**C**) Quantification of the extent of LY6A protein reduction in cultures treated with no sgRNA (–), or sgRNAs targeted to *Ly6a or Ly6c1*.

**Supplementary Table 1.**
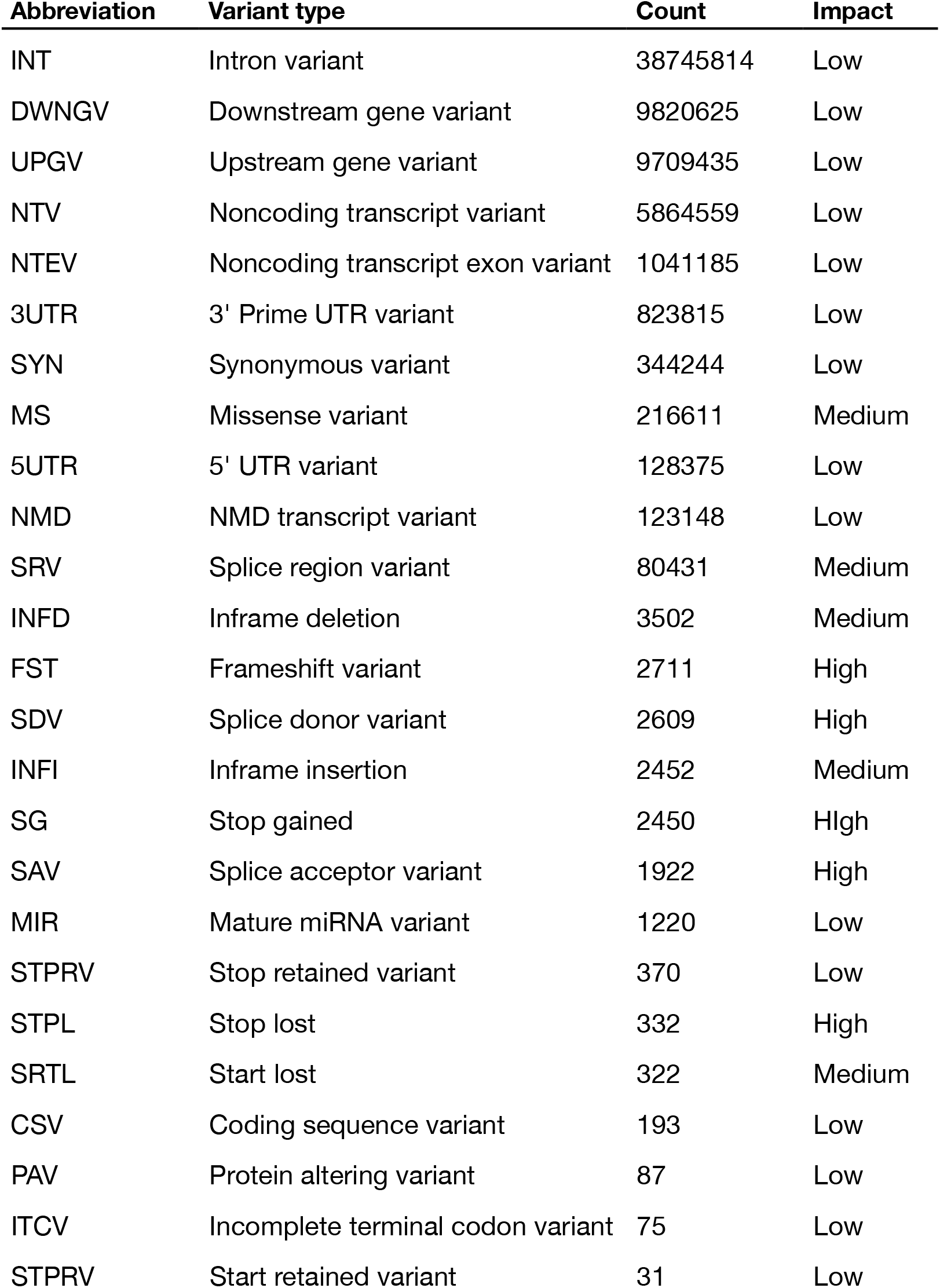
The types of genetic variants included in the linkage study. The predicted variant types, their count among all 36 mouse strains in the in the mouse genome project^22^ database, and their predicted impact is shown. We restricted the analysis to variants with medium or high predicted impact on gene expression or coding sequence.

| Abbreviation | Variant type | Count | Impact |
| --- | --- | --- | --- |
| INT | Intron variant | 38745814 | Low |
| DWNGV | Downstream gene variant | 9820625 | Low |
| UPGV | Upstream gene variant | 9709435 | Low |
| NTV | Noncoding transcript variant | 5864559 | Low |
| NTEV | Noncoding transcript exon variant | 1041185 | Low |
| 3UTR | 3' Prime UTR variant | 823815 | Low |
| SYN | Synonymous variant | 344244 | Low |
| MS | Missense variant | 216611 | Medium |
| 5UTR | 5' UTR variant | 128375 | Low |
| NMD | NMD transcript variant | 123148 | Low |
| SRV | Splice region variant | 80431 | Medium |
| INFD | Inframe deletion | 3502 | Medium |
| FST | Frameshift variant | 2711 | High |
| SDV | Splice donor variant | 2609 | High |
| INFI | Inframe insertion | 2452 | Medium |
| SG | Stop gained | 2450 | High |
| SAV | Splice acceptor variant | 1922 | High |
| MIR | Mature miRNA variant | 1220 | Low |
| STPRV | Stop retained variant | 370 | Low |
| STPL | Stop lost | 332 | High |
| SRTL | Start lost | 322 | Medium |
| CSV | Coding sequence variant | 193 | Low |
| PAV | Protein altering variant | 87 | Low |
| ITCV | Incomplete terminal codon variant | 75 | Low |
| STPRV | Start retained variant | 31 | Low |

**Supplementary Table 2.**
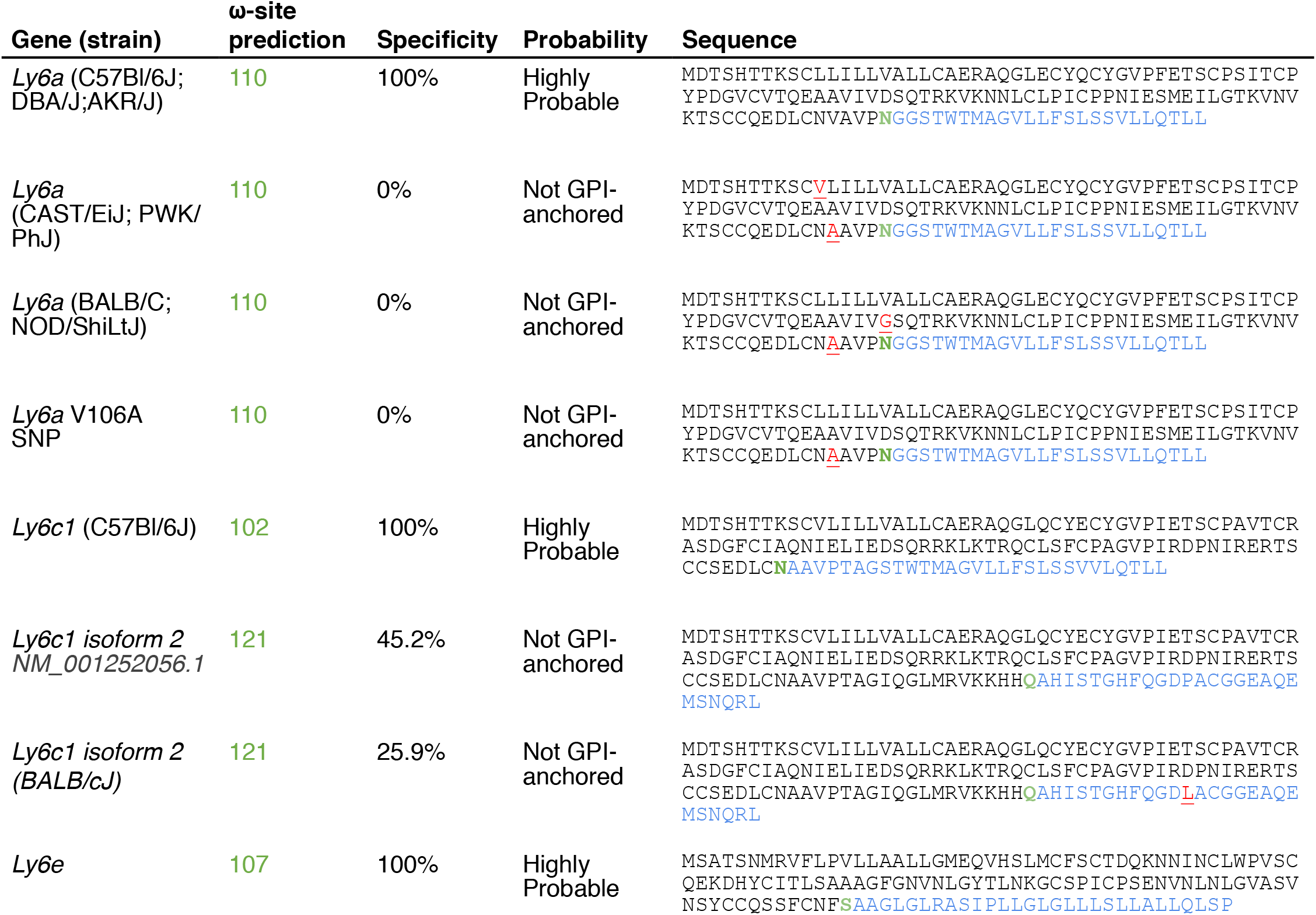
LY6A from C57Bl/6J but not BALB/cJ mice is predicted to be GPI anchored. The predGPI prediction model^33^ was used to assess the probability that the alleles of *Ly6a* and *Ly6c1* present in permissive and nonpermissive mouse strains encode protein products likely to be modified by a GPI anchor. The Ly6c1 missense SNP affects splice isoform 2, but not isoform 1. Within the amino acid sequences, SNPs are highlighted in red and underlined text; predicted GPI-anchor sites (ω-sites) are highlighted in bold green text, and cleaved C-terminal hydrophobic tail sequences are highlighted in blue text.

**Supplementary Table 3.** Permissive or nonpermissive AAV-PHP.eB CNS transduction phenotypes for inbred mouse lines with available WGS data. The lines highlighted in bolded text were used in the analysis presented in Supplementary Fig. 2A.

| Strain | Enhanced CNS tropism | References |
| --- | --- | --- |
| **AKR/J** | Present | This study |
| **BTBR/T+/Itpr3tf/J** | Present | Predicted |
| **C57BL/6J** | Present, previously reported | 5,6,9,19 |
| **C57BL/6NJ** | Present, previously reported | 10 |
| **C57BL/10J** | Present | Predicted |
| **C57BR/cdJ** | Present | Predicted |
| **C57L/J** | Present | This study |
| C58/J | Present | Predicted |
| **DBA/1J** | Present | Predicted |
| **DBA/2J** | Present | This study |
| **FVB/NJ** | Present | This study |
| I/LnJ | Present | Predicted |
| KK/HiJ | Present | Predicted |
| **LP/J** | Present | This study |
| **NZW/LacJ** | Present | Predicted |
| RF/J | Present | Predicted |
| **MOLF/EiJ** | Present | This study |
| 129P2/OlaHsd | Present | Predicted |
| **129S1/SvImJ** | Present | Predicted |
| 129S5SvEvBrd | Present | Predicted |
| **A/J** | Absent | Predicted |
| **BALB/cJ** | Absent, previously reported | <sup>16</sup>; this study |
| BUB/BnJ | Absent | Predicted |
| **CAST/EiJ** | Absent | This study |
| **CBA/J** | Absent | This study |
| C3H/HeH | Absent | Predicted |
| **C3H/HeJ** | Absent | Predicted |
| **LEWES/EiJ** | Absent | Predicted |
| **NOD/ShiLtJ** | Absent | This study |
| **NZB/B1NJ** | Absent | This study |
| **NZO/HILtJ** | Absent | Predicted |
| **PWK/PhJ** | Absent | This study |
| SEA/GnJ | Absent | Predicted |
| **SPRET/EiJ** | Absent | Predicted |
| SEA/GnJ | Absent | Predicted |
| ST/bJ | Absent | Predicted |
| **WSB/EiJ** | Absent | Predicted |
| ZALENDE/EiJ | Absent | Predicted |

